# Pro-inflammatory role of granzyme K producing bystander CD8^+^ T cells in acute myeloid leukemia

**DOI:** 10.1101/2025.08.12.669682

**Authors:** Lisa Aziez, Nicolas Deredec, Ismael Boussaid, Carolyn G. Shasha, Romain Vazquez, Chloé Friedrich, Ania Alik, Kanchanadevi Manasse, Zoé Fremont-Debaene, Alexandra Barthelemy, Cyril Catelain, Philippe Rameau, Marguerite Vignon, Justine Decroocq, Olivier Kosmider, Dorothée Selimoglu-Buet, Evan W. Newell, Eric Solary, François Delhommeau, Olivier Herault, Eric Tartour, Rudy Birsen, Didier Bouscary, Michaela Fontenay, Nicolas Chapuis, Yannick Simoni

## Abstract

Acute myeloid leukemia (AML) is a heterogeneous group of blood malignancies with a 5-year survival rate below 30%, highlighting the urgent need for more effective therapeutic strategies. T cell-based immunotherapies have demonstrated remarkable success in solid tumors, yet the role of CD8^+^ T cells in AML remains unclear. In this study, we analyzed the composition, antigenic specificity, and function of CD8^+^ T cells in paired blood and bone marrow samples from AML patients. While we did not identify exhausted CD8^+^ T cells as seen in solid tumors, we observed a distinct population of functional CD69^+^ CD8^+^ T cells specifically enriched in the bone marrow. These cells primarily recognized non-tumor antigens, including epitopes derived from Epstein–Barr virus (EBV) and cytomegalovirus (CMV). Notably, this bystander CD8^+^ T cell population showed high expression of Granzyme K, a cytokine found in the bone marrow of AML patients. Granzyme K did not induce leukemic cell death but instead promoted the secretion of IL-8, a pro-inflammatory cytokine known to play a detrimental role in AML pathology. Rather than mounting an anti-tumor response, these CD8^+^ T cells contribute to a pro-inflammatory environment that may exacerbate AML progression and severity. These findings provide a rationale for exploring therapeutic strategies aimed at inhibiting pro-inflammatory CD8^+^ T cells and targeting Granzyme K activity in association with actual therapies.

**Graphical Abstract:** 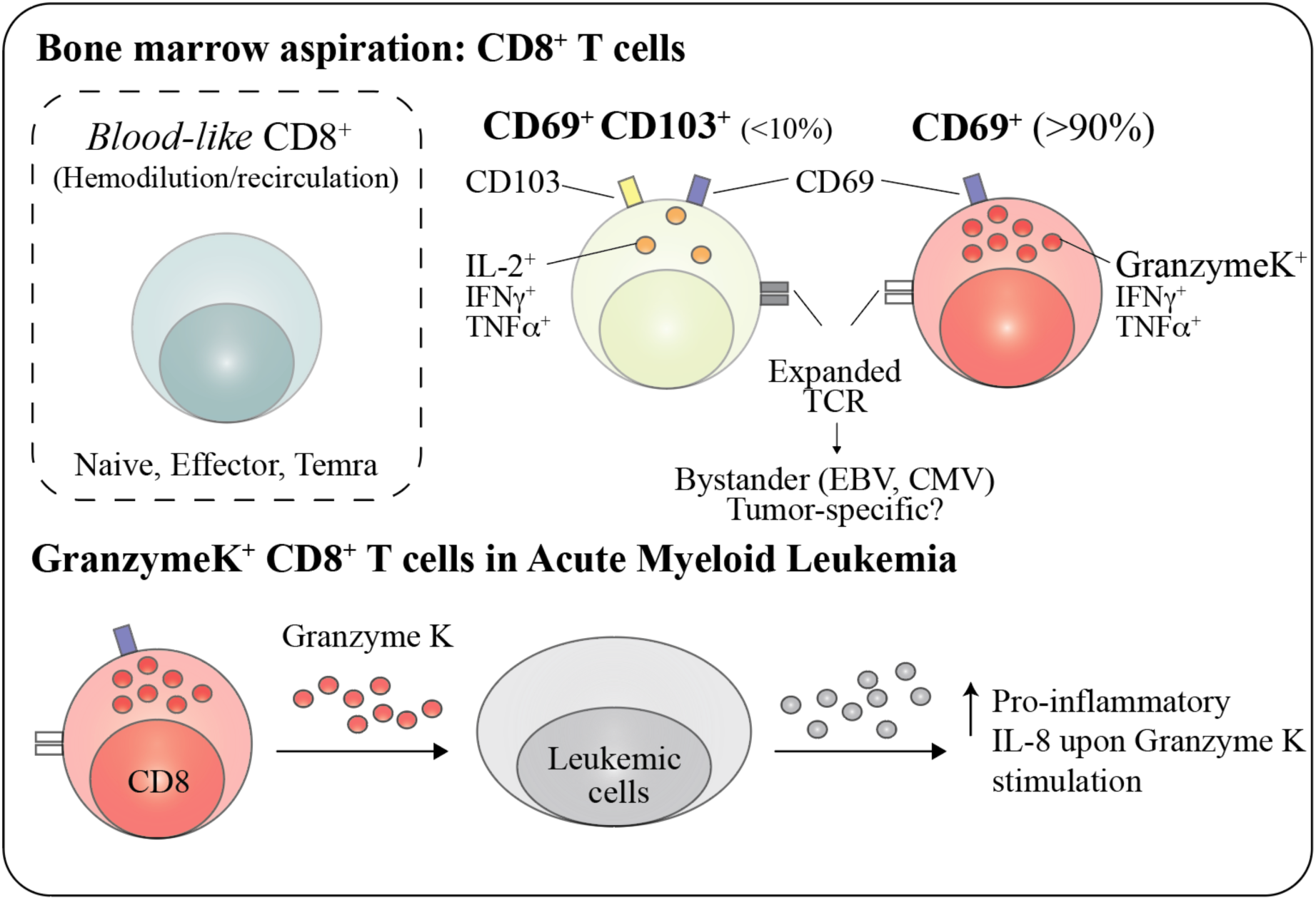

## Introduction

Acute myeloid leukemia (AML) is a heterogeneous group of blood malignancies characterized by the uncontrolled proliferation of bone marrow derived immature myeloid progenitors. Despite advances in treatment, prognosis remains poor, with a 5-year survival rate of approximately 30–40% for patients under 60 and less than 20% for those over 60^1^. This underscores the urgent need for more effective therapeutic strategies. The bone marrow immune microenvironment plays a critical role in AML pathophysiology, such as T lymphocytes, engaging in complex interactions with leukemic cells^2^. To evade immune surveillance, AML cells employ multiple mechanisms, including downregulation of major histocompatibility complex (MHC) molecules, upregulation of inhibitory ligands, reduced expression of activating ligands, and modulation of soluble factors in the microenvironment^3^. In recent years, immunotherapeutic strategies targeting T cells have transformed cancer treatment. In solid tumors, immune checkpoint inhibitors such as anti-PD-1 and anti-CTLA-4 therapies can reinvigorate tumor-specific exhausted T cells to kill tumor cells^4^. In hematologic malignancies, bispecific antibodies (BsAbs), such as CD3xBCMA^5^, or chimeric antigen receptor (CAR) T-cell therapies^6^ have been developed to enhance T-cell cytotoxicity against tumor cells. However, these approaches have shown limited success in AML^7^. Although reports suggest that CD8^+^ T cells in the AML bone marrow microenvironment may exhibit exhaustion or dysfunction, traducing the presence of tumor-specific T cells^8–10^, conflicting evidence indicates the presence of fully functional CD8^+^ T cells in AML^10,11^. To address this discrepancy and investigate the contribution of CD8^+^ T cells in AML pathology, we used high-throughput approaches, such as mass-cytometry (CyTOF), Cellular Indexing of Transcriptomes and Epitopes by Sequencing (CITE-seq) and functional assay, to characterize the composition, antigenic specificity, and functional properties of CD8^+^ T cells in blood and bone marrow samples from AML patients.

## Results

### CD8^+^ T cells in AML bone marrow lack classical exhaustion markers

Using mass cytometry and a dedicated panel for in-depth profiling, we analyzed CD8^+^ T cells from bone marrow aspirates of AML patients (Table 1 and Figure 1A). In order to rule out contamination by leukemic cells expressing aberrant lymphoid markers, leukemic derived myeloid cells were gated out (CD33^+^ CD13^+^) among live immune cells (DNA^+^ Cisplatin^-^ CD45^+^). Among T cells (CD3^+^), Tψο (TCRψο^+^) and MAIT (TCR Vα7.2^+^ CD161^+^) cells were excluded to identify conventional Tαβ CD8^+^ T cells, representing 4.7% among CD45^+^ cells in AML bone marrow (Supplementary Figure 1A). As observed in solid cancer^12^, CD8^+^ T cells are highly heterogeneous population among AML patient, with uniform manifold approximation and projection (UMAP) visualization showing multiple distinct cells clusters across patients (Figure 1B). However, unlike in solid tumors, we did not observe a distinct cluster of exhausted CD8^+^ T cells^13–15^, a hallmark of chronically stimulated tumor-specific T cells (Figure 1C, Supplementary Figure 1B). To further investigate immune checkpoint expression, we compared inhibitory receptor profiles of AML’s bone marrow CD8^+^ T cells with tumor-infiltrating lymphocytes from colorectal, lung, and kidney cancers from public dataset^12,16^. AML-associated CD8^+^ T cells exhibited low expression of TIM-3 (1.2%) and TIGIT (3.6%), along with significantly reduced levels of PD-1 (22.3%) and CD39 (2.9%) (Figure 1D). Co-expression of multiple inhibitory receptors, a defining feature of T cell exhaustion a hallmark of tumor-specific T cells^12,17^ ^18^, was rare (Figure 1E). PD-1 and CD39 were co-expressed in only 0.8% of CD8^+^ T cells, a frequency markedly lower than in solid tumors (Figure 1F). Interestingly, only a small subset (2.7%) of bone marrow CD8^+^ T cells co-expressed CD69 and CD103, a phenotype characteristic of a canonical tissue-resident memory T cells (Trm)^19^. However, a much larger proportion (22%) expressed CD69 alone, a marker associated with tissue-residency and recently activated T cells, that has been reported in other hematological malignancies ^20^. These frequencies were significantly lower than those observed in solid tumors (Figure 1F). Among CD8^+^ T cells lacking CD69, we identified subsets resembling circulating blood T cells, including a terminally differentiated effector memory (Temra-like) population expressing CD57^+/−^, KLRG1^+^, CD28^−^, and CD45RO^−^, as well as an effector memory-like (Effector-like) population expressing CD45RO^+^, CD28^+^, and CD127^+^. Naïve-like CD8^+^ T cells (Naïve-like) population expressing CD45RO^-^ CCR7^+^, typically found in circulation, were also present in bone marrow samples (Supplementary Figure 1C). Collectively, our findings suggest that CD8^+^ T cells in AML bone marrow aspirates lack conventional exhaustion signatures observed in solid tumors. Instead, these cells exhibit phenotypic diversity, including subsets expressing tissue residency markers and others resembling circulating blood T cells.

**Figure 1.**
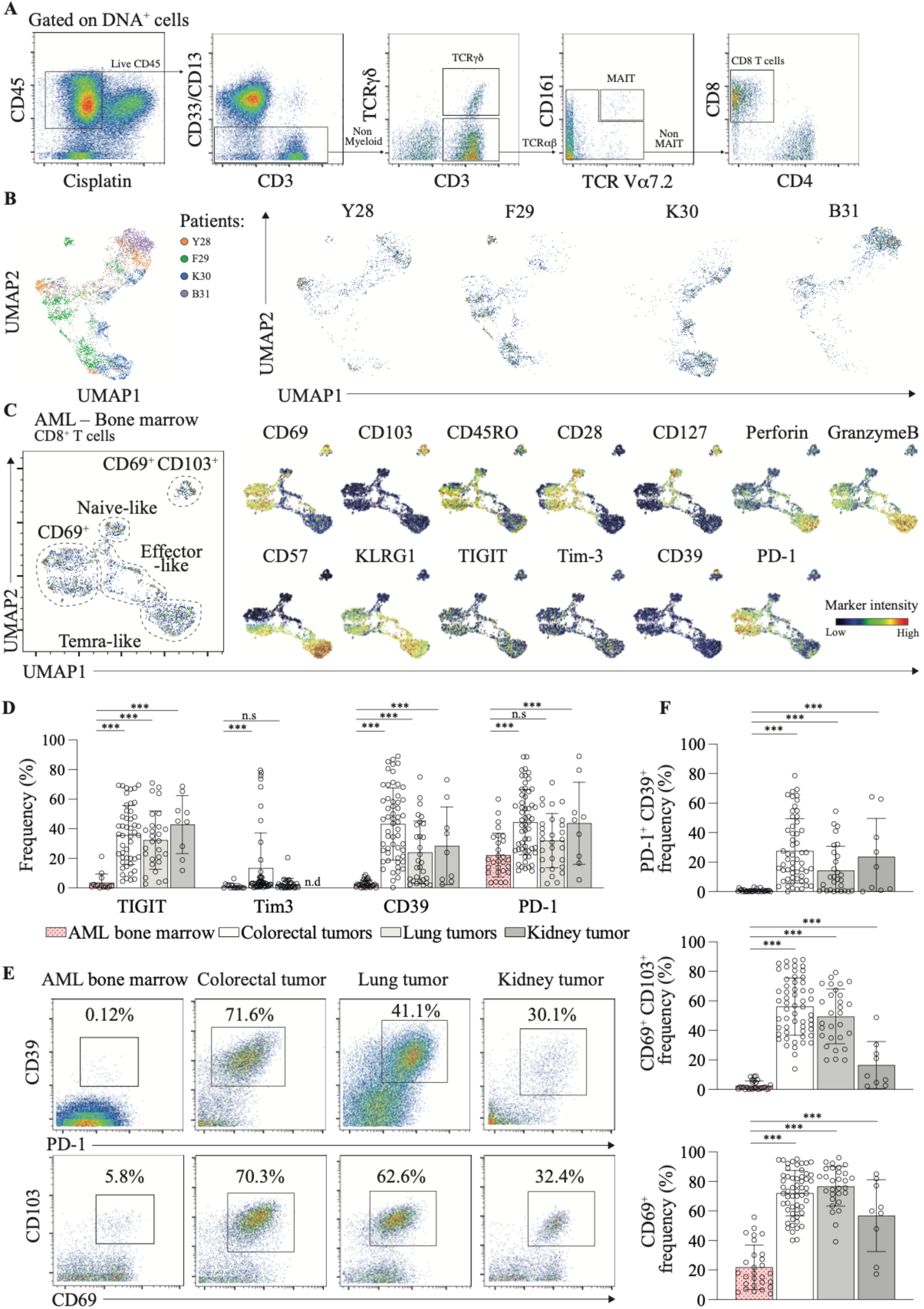
CD8^+^ T cells in bone marrow samples from AML patients do not show explicit signs of exhaustion. **A**. Gating strategy used to identify CD8⁺ T cells from bone marrow aspirates of an AML patient (gated on DNA^+^ cells). Representative data from one patient. Mass cytometry analysis **B**. UMAP analysis of CD8⁺ T cells from AML bone marrow: combined view (left panel) colored by patient and individual projections (right panel) for Y28, F29, K30, and B31. **C**. UMAP plot of CD8^+^ T cells from bone marrow aspirates of an AML patient, analyzed by mass cytometry (left panel). Normalized marker expression intensities were calculated and overlaid on the UMAP plot, allowing the identification of naïve-like, effector-like, Temra-like, CD69^+^ T cells and CD69^+^ CD103^+^ CD8^+^ T cells (right panel). Representative data from one patient (F29). Mass cytometry data. **D.** Expression of selected co-inhibitory markers, hallmarks of exhaustion, by CD8^+^ T cells from AML bone marrow (n=14 to 28 patients), colorectal tumors (n=48 to 56 patients), lung tumors (n=29), and kidney tumors (n=9). **E.** Mass cytometry dot plots representing the co-expression of PD-1 and CD39 (top panel) and co-expression of CD69 and CD103 (bottom panel). **F.** Representative expression of the co-expression of PD-1 and CD39 (top panel), co-expression of CD69 and CD103 (middle panel) and expression of CD69 (bottom panel) by CD8^+^ T cells from AML bone marrow (n=31), colorectal tumors (n=56), lung tumors (n=29), and kidney tumors (n=9). All data are from biologically independent individuals. Means ± SD, Mann-Whitney test, two-tailed. *p ≤ 0.05, **p ≤ 0.01, ***p ≤ 0.001.

### CD69^-^ CD8^+^ T cell populations in AML bone marrow aspirates reassemble circulating blood T cells

The potential contamination of bone marrow aspirates by peripheral blood, or hemodilution, has been previously reported in the context of hematologic malignancy diagnosis^21–24^. To assess whether CD8^+^ T cells lacking CD69 expression represent *blood-derived* cells, we analyzed phenotypic changes between paired peripheral blood mononuclear cells (PBMCs) and bone marrow aspirates samples from AML patients (Figure 2A). UMAP analysis revealed significant overlap between bone marrow and PBMC derived CD8^+^ T cells. These *blood-like* CD8^+^ T cells accounted for more than 70% of bone marrow CD8^+^ T cells in our cohort (Figure 2B). Notably, while a similar population is observed in solid tumors, it represents only a minor fraction of tumor-infiltrating CD8^+^ T cells compared to AML (Figure 2B and Supplementary Figure 2A). The *blood-like* CD8^+^ T cell population in AML bone marrow comprised about 30% Naïve-like, 55% Effector-like, and 29% Temra-like CD8^+^ T cells (Figure 2C). To confirm this observation, we performed single-cell TCRαβ sequencing using CITE-seq on sorted CD8^+^ T cells from paired PBMC and bone marrow samples (Supplementary Figure 2B). The Naïve-like subset exhibited 99.9% unique TCR clonotypes, with transcriptomic profiles indistinguishable from their blood-derived counterparts (Figures 2D, E, G and Supplementary Figures 3A and C). Expanded TCR clonotypes, a hallmark of clonal expansion of antigen-experienced T cells, were observed within the Effector-like and Temra-like subsets. These clonotypes were highly similar between blood and bone marrow (Figure 2F), with correlated frequencies and similar spatial distributions in UMAP protein dataset projections (Figures 2G, H, and Supplementary Figures 3B and D). These findings indicate that a significant fraction of CD8^+^ T cells in AML bone marrow aspirates closely resembles circulating blood T cells, suggesting they originate from peripheral blood rather than being specific to the bone marrow microenvironment.

**Figure 2.**
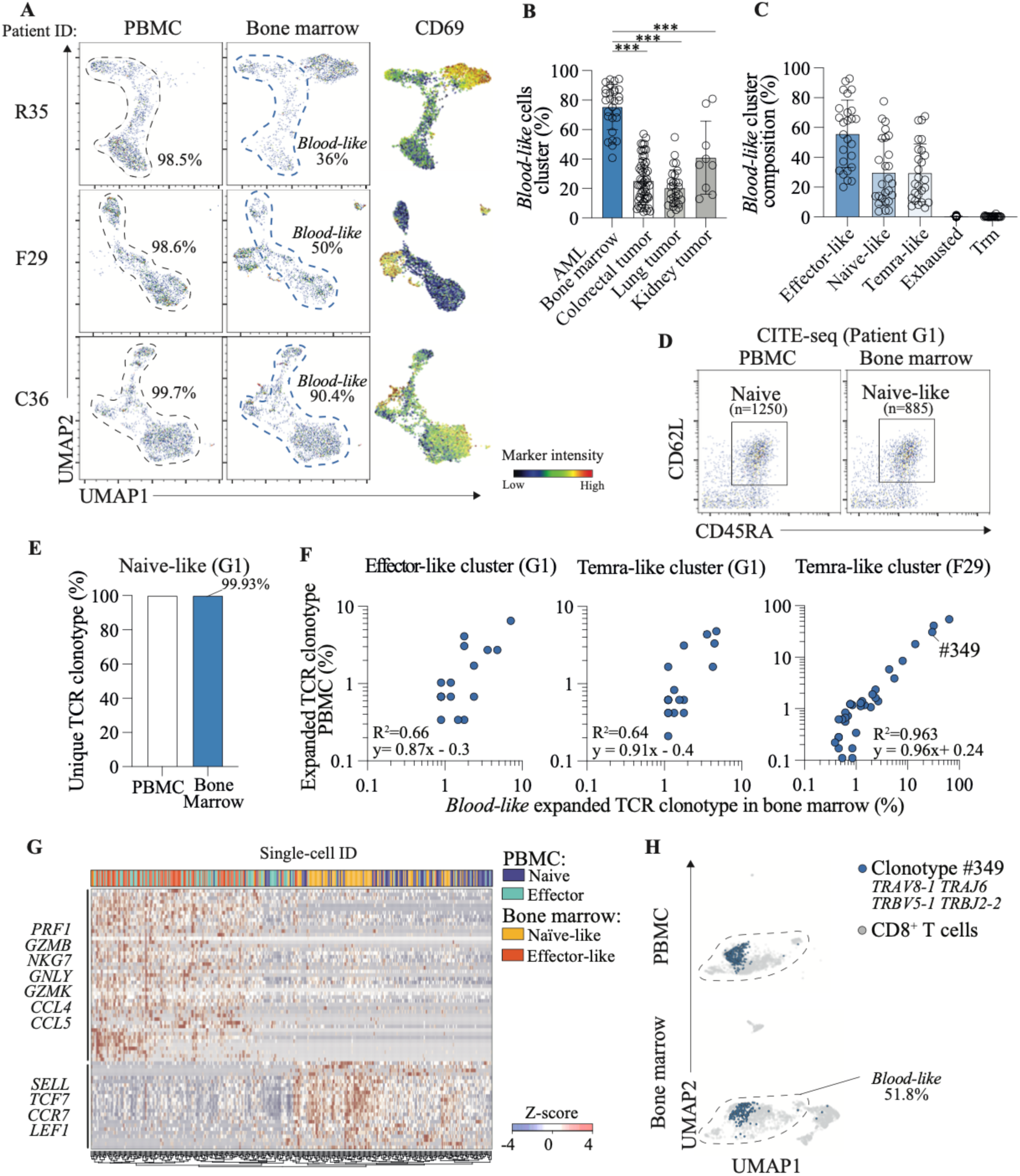
CD69^-^ CD8^+^ T Cell populations in bone marrow samples display similarities with their blood counterpart. **A.** Parallel mass cytometry analysis of CD8^+^ T cells from PBMCs and bone marrow aspirates of AML patients, allowing for the identification of the *blood-like* CD8**^+^** T cell cluster (light blue). UMAP was performed separately for each patient (left panel). Normalized expression intensities of CD69 were calculated and overlaid on the UMAP plot (right panel). **B.** Quantification of the *blood-like* cell cluster among CD8^+^ T cells from AML bone marrow aspirates (n=28), colorectal tumors (n=56), lung tumors (n=29), and kidney tumors (n=9). Data represent biologically independent individuals. Means ± SD, Mann-Whitney test, two-tailed. *p ≤ 0.05, **p ≤ 0.01, ***p ≤ 0.001. **C.** Quantification of effector-like, naïve-like, Temra-like, exhausted and Trm CD8**^+^** T cells within the *blood-like* cell cluster from AML bone marrow (n=26 to 27 patients) by mass cytometry. Data are from biologically independent individuals. **D.** CITE-seq dot plots showing CD8^+^ T cells from PBMCs and bone marrow for the identification of naïve T cells using CD62L and CD45RO markers. Data representative of one patient (G1). **E.** Frequency of unique TCR clonotypes, defined by unique V(D)J nucleotide sequences for the paired alpha and beta chains, measured by CITE-seq, among naïve-like cells in blood (n=1250 single cells) and bone marrow (n=885 single cells). Data representative of one patient (G1). **F.** Scatterplots comparing the frequency of expanded TCR clonotypes (≥5 cells), measured by CITE-seq, in paired PBMC and bone marrow samples within the *blood-like* cluster, separated by patient and population (effector-and Temra-like cells). R^2^ = coefficient of determination, y = linear regression equation. **G**. Heatmap shows Z-scores of the average expression of differentially expressed genes measured by CITE-seq from naïve and effector-like CD8^+^ T cells in paired PBMC and bone marrow samples (patient G1). Data represent a sample of 100 single cells per cluster. **H.** Projection of expanded TCR clonotype #349 in PBMC (upper panel) and BM (lower panel) on the UMAP plot generated from the surface marker dataset from CITE-seq.

### CD69^+^ CD8^+^ T cells in AML bone marrow are tissue-specific and express Granzyme K

In contrast to the *blood-like* CD8^+^ T cells, distinct populations of CD8^+^ T cells expressing CD69 were found to be bone marrow tissue-specific. UMAP analysis of paired tissues confirmed that CD69^+^ CD8^+^ T cells were restricted to the bone marrow (Figure 3A). Among these, fewer than 10% expressed CD103, CD49a, CD127, or CD101, markers typically associated with Trm cells^25^. Of note, around 30% of the CD69^+^ CD103^+^ CD8^+^ T cells express CD39, a marker associated with tumor-specific cells, but lack PD-1, TIGIT, Tim-3, CTLA-4 expression (Figure 3B and C). Instead, the majority of bone marrow specific CD8^+^ T cells (90%, CD69^+^ CD103^-^) exhibited a non-exhausted, non-senescent phenotype characterized by CD28, KLRG1, and PD-1 expression, consistent with long-lived effector memory cells^26^ (Figures 3B and C).

**Figure 3.**
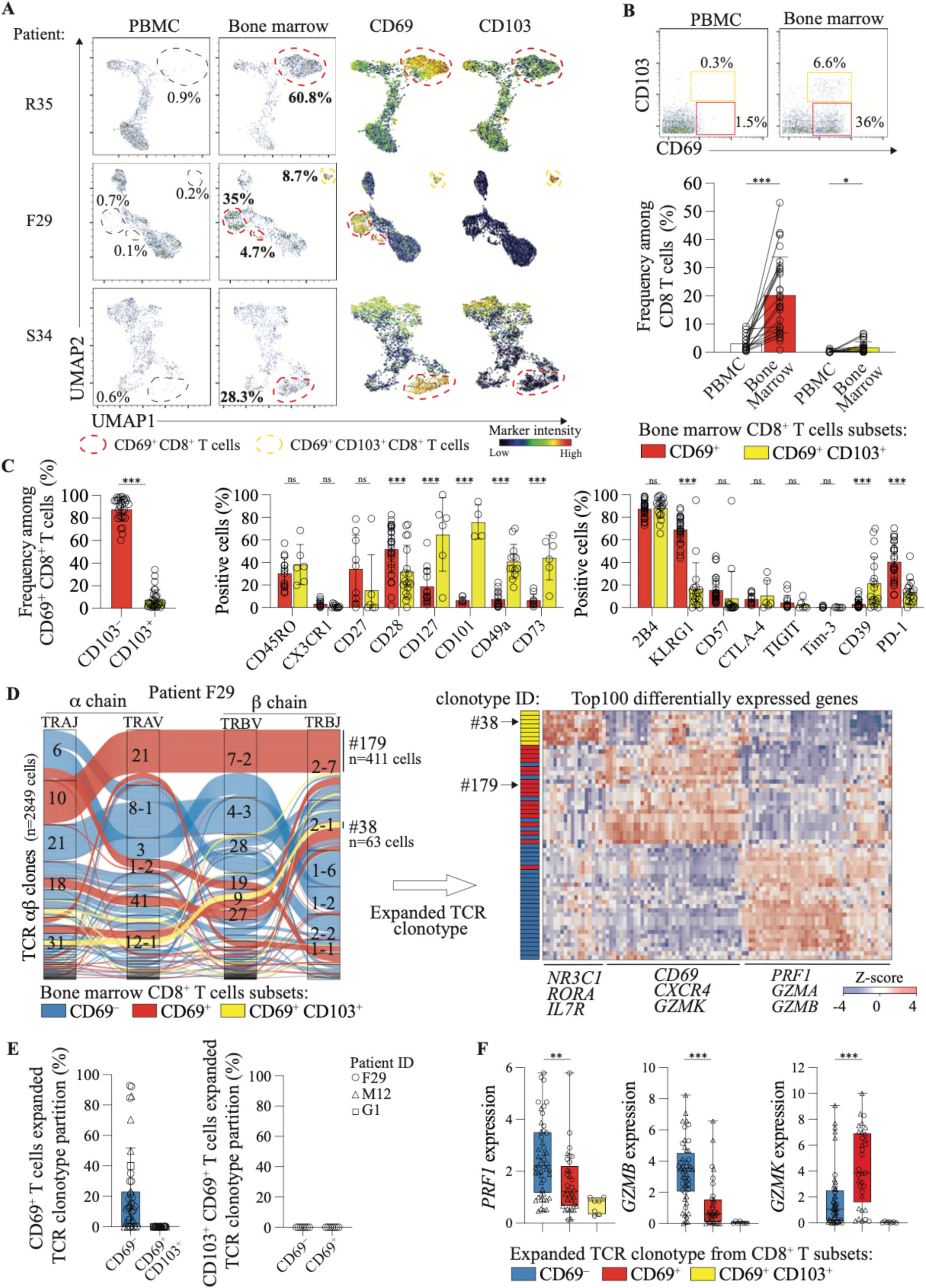
CD69^+^ CD8^+^ T cells are specific to bone marrow tissue. **A**. Parallel mass cytometry analysis of CD8^+^ T cells from PBMCs and bone marrow aspirates of AML patients, allowing for the identification of bone marrow-specific CD69^+^ (red) and CD69^+^ CD103^+^ (yellow) CD8^+^ T cells. UMAP was performed separately for each patient (left panel). Normalized expression intensities of CD69 and CD103 were calculated and overlaid on the UMAP plot (right panel). **B.** Mass cytometry dot plots illustrating the expression of CD69^+^ (red) and CD69^+^ CD103^+^ (yellow) by CD8^+^ T cells in PBMCs and bone marrow are presented. Representative data from one AML patient are shown (upper panel). Frequency of CD69^+^ and CD69^+^ CD103^+^ CD8^+^ T cells in PBMCs and bone marrow of AML patients (bottom panel) (n= 17 pairs of patients) Means ± SD, with statistical significance assessed using a paired t-test, two-tailed. *p ≤ 0.05, **p ≤ 0.01, ***p ≤ 0.001. **C.** Frequency of CD69^+^ (red, n = 30 patients) and CD69^+^ CD103^+^ (yellow, n = 28 patients) CD8^+^ T cells among the bone marrow specific population of AML patients (left panel). Expression of selected activation (middle panel) and inhibitory (left panel) markers by bone marrow specific CD69^+^ and CD69^+^ CD103^+^ CD8^+^ T cells. Means ± SD, with statistical significance assessed using a paired t-test, two-tailed. *p ≤ 0.05, **p ≤ 0.01, ***p ≤ 0.001. **D**. Alluvial plot displaying the TCR clone composition measured by CITE-seq of CD69^-^ *blood-like* cluster (blue), and CD69^+^ (red), CD69^+^ CD103^+^ (yellow) bone marrow specific clusters detected in patient F29 (middle panel). Distribution of the top expanded TCR clonotypes (detected in ≥ 5 cells) among CD8^+^ T cells subsets. Heatmap comparing differentially expressed genes by expanded TCR clonotypes detected in CD69^+^ and CD69^-^ CD8^+^ T cells from bone marrow is presented, as measured by CITE-seq (right panel). **E**. Partition of expanded TCR clonotype from CD69^+^ CD8^+^ T cells (left panel) and CD103^+^ CD69^+^ CD8^+^ T cells (right), among each CD8^+^ T cells subsets. Expanded TCR from Patient F29 (Circle), patient M12 (triangle) and patients G1 (square). **F** Expression of cytotoxic gene *PRF1* (perforin), *GZMB* (granzyme B) and *GZMK* (granzyme K) from expanded TCR clonotype from CD8^+^ T subsets: CD69^-^ (blue), CD69^+^ (red), CD69^+^ CD103^+^ (yellow).

Single-cell TCR analysis revealed that CD69⁺ CD8⁺ T cells exhibit a distinct expanded TCR repertoire compared to the *blood-like* CD69⁻ CD8⁺ T cell population (Figure 3D, Supplementary Figure 4). However, the presence of shared expanded TCR clonotypes between CD69^-^ and CD69⁺ CD8^+^ T cells implies that they may, at least in part, recognize similar antigens (Figure 3E, Supplementary Figure 4). Moreover, CD69⁺ CD103⁺ subset displayed non-overlapping TCR clonotypes, confirming that they are distinct populations with differing antigen specificities (Figure 3E). At the transcriptomic level, CD69^+^ CD8^+^ T cells exhibited a unique gene signature characterized by high expression of *GZMK* (granzyme K), while lacking *GZMB* (granzyme B) and *PRF1* (perforin), which are typically associated with cytotoxic T cells (Figures 3D, F and Supplementary Figures 5, 6 and 7).

### Granzyme K⁺ bystander CD8⁺ T cells are enriched in AML bone marrow

*In-silico* analysis of antigenic specificity using public TCR database revealed the presence of an expanded TCR clonotype specific for Epstein-Barr virus (EBV) peptides within the CD69⁺ CD8⁺ T cell population, suggesting the presence of bystander T cells (Figure 4A). To validate this finding at the protein level, antigen specificity was assessed using MHC class I tetramers loaded with viral antigens from EBV, cytomegalovirus (HCMV), Influenza, and SARS-CoV-2 (Table 2, Supplementary Figure 8). This confirmed the presence of cancer-unrelated, EBV-or HCMV-specific CD69⁺ CD8⁺ T cells in the bone marrow (Figures 4B, C, Supplementary Figure 8A, B and C). However, no cancer-unrelated in CD69^+^ CD103^+^ CD8^+^ T were detectable in our cohort (Supplementary Figure 8D). Functional assays demonstrated that CD69⁺ and CD69⁺ CD103⁺ CD8⁺ T cells are functionally competent and not exhausted, as indicated by their ability to produce IFNγ and TNFα upon stimulation. Notably, CD69⁺ CD103⁺ CD8⁺ T cells also produced IL-2, a cytokine implicated in T cells growth, proliferation, and survival (Figure 4D). In terms of cytotoxic potential, both CD69⁺ and CD69⁺ CD103⁺ CD8⁺ T cells exhibited limited cytotoxicity, with fewer than 20% expressing Perforin or Granzyme B. In contrast, over 60% of CD69⁺ CD8⁺ T cells expressed Granzyme K in AML bone marrow (Figure 4E). Collectively, our findings demonstrate that bone marrow CD69⁺ CD8⁺ T cells are specific for cancer-unrelated antigens and display a distinct transcriptomic, phenotypic and functional signature characterized by the expression of Granzyme K.

**Figure 4.**
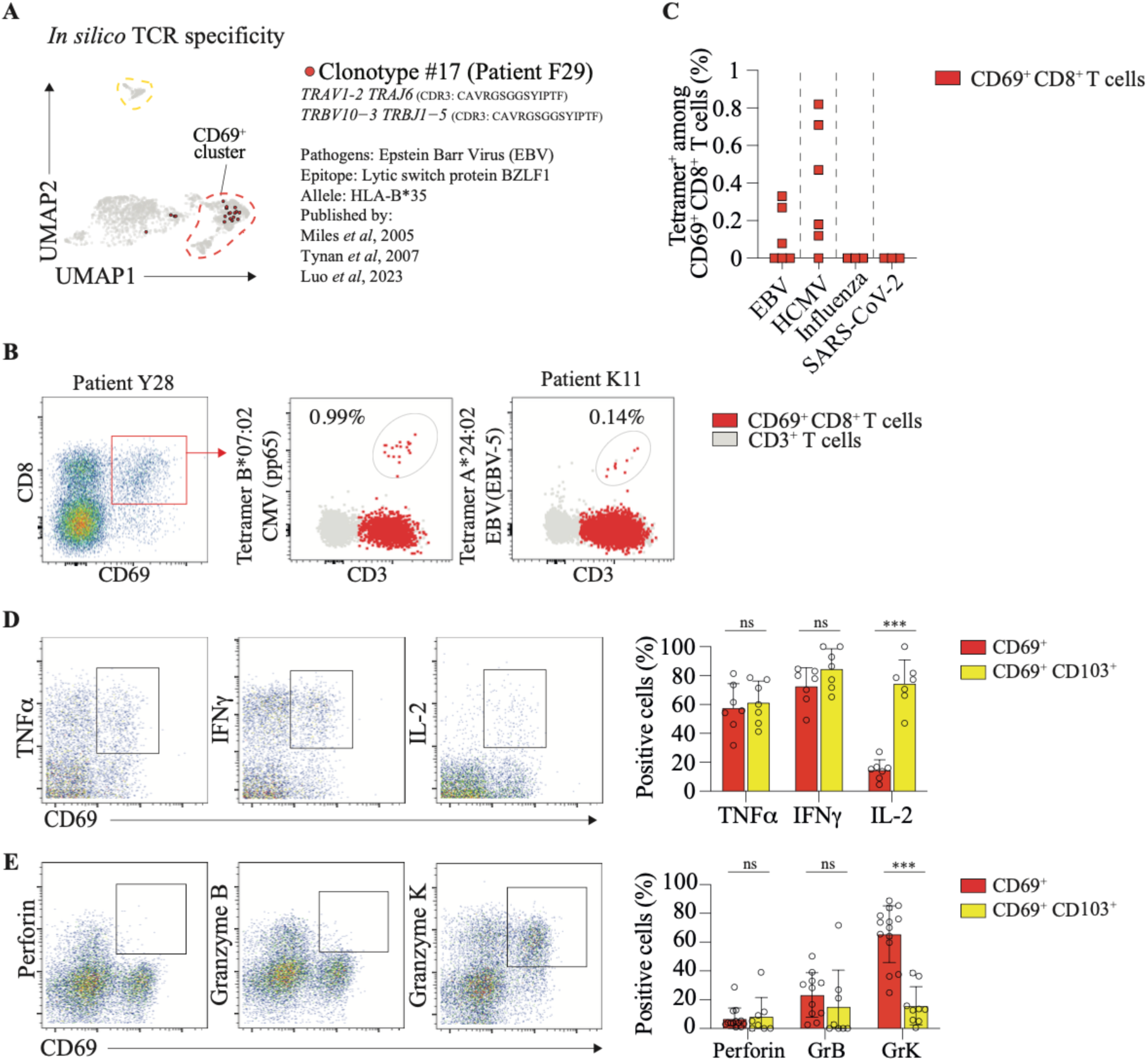
Identification of Granzyme. **K**^+^ **bystander CD8**^+^ **T Cells in Bone marrow of AML Patients. A.** *In silico* analysis revealed that the CD69^+^ expanded TCR clonotype #17 (from AML patient F29) matched sequences in public databases, indicating its antigenic specificity. **B.** Representative flow cytometry dot plot showing cancer-unrelated CD8^+^ T cells specific for different epitopes, identified in bone marrow specific CD69^+^ CD8^+^ T cells (red) of AML patients using mass cytometry screening. Percentages are of MHC tetramer^+^ cells among CD8^+^ T cells for each patient. **C.** Frequencies of cancer-unrelated CD8^+^ T cells identified by mass cytometry in CD69^+^ CD8^+^ T cells from bone marrow. **D.** Mass cytometry dot plots illustrating the expression of IFNγ, TNFα, and IL-2 in AML bone marrow (left panel) and their expression frequency measured by mass-cytometry in CD69^+^ (red) and CD69^+^ CD103^+^ (yellow) CD8^+^ T cells stimulated with PMA/ionomycin and Brefeldin A for 4h (right panel) (n = 7). **E.** Mass cytometry dot plots illustrating the expression of Perforin, Granzyme B, and K in relation to CD69 in AML bone marrow (left panel) and their expression frequency measured by mass-cytometry in CD69^+^ (red) and CD69^+^ CD103^+^ (yellow) CD8^+^ T cells (n = 11 to 14 patients). Means ± SD, multiple paired t-test, two-tailed. *p ≤ 0.05, **p ≤ 0.01, ***p ≤ 0.001.

### Granzyme K-producing CD8^+^ T cells contribute to a pro-inflammatory environment in AML

Given that Granzyme K-expressing CD8⁺ T cells have been described in cancer-unrelated contexts, such as immune aging, we investigated whether these cells were specifically induced by the AML microenvironment or were also present in the bone marrow of healthy individuals. Mass cytometry analysis revealed a similar population of CD8⁺ T cells, expressing CD69 and Granzyme K but lacking Perforin and Granzyme B, in the bone marrow of healthy donors (Figure 5A). UMAP analysis of CITE-seq data confirmed that the two populations shared a similar phenotype (Supplementary Figures 8E, F and G). At the transcriptomic level, both populations displayed a comparable gene expression signature, confirming their similarity (Figure 5B). We next quantified Granzyme K levels in the bone marrow microenvironment of healthy donors and AML patients. Quantitative analysis revealed significantly elevated concentrations of Granzyme K in AML medullary plasma (mean = 932.2 pg/mL) compared to age-matched healthy donors (mean = 103.3 pg/mL). Interestingly, Granzyme K was barely detectable in the medullary plasma of approximately 30% of AML patients analyzed (<10 pg/mL) (Figure 5C). To assess the functional impact of Granzyme K on leukemic cells, AML cell lines were treated with recombinant Granzyme K. This treatment had no effect on cell viability (Figure 5D). However, Granzyme K significantly enhanced IL-8 secretion, but only in AML cell lines with constitutive IL-8 expression. No significant changes were observed in the secretion of IL-1β, IL-6, TNFα, or S100A8/A9 (Figure 5E, Supplementary Figures 9A and B). Similar findings were observed in primary leukemic cells from patients where Granzyme K did not induce cell death but did enhance IL-8 production (Figures 5F, G, and H, Supplementary Figures 9C and D). As with the AML cell lines, IL-8 secretion in response to Granzyme K was heterogeneous across patient samples. Together, these findings indicate that the presence of Granzyme K⁺ CD69⁺ CD8⁺ T cells in the bone marrow microenvironment is not specific to AML patients. However, their activation and subsequent release of Granzyme K appear to be restricted to the AML context. The production of Granzyme K by these cells contributes to increase a pro-inflammatory microenvironment, which may in turn exacerbate AML progression and severity.

**Figure 5.**
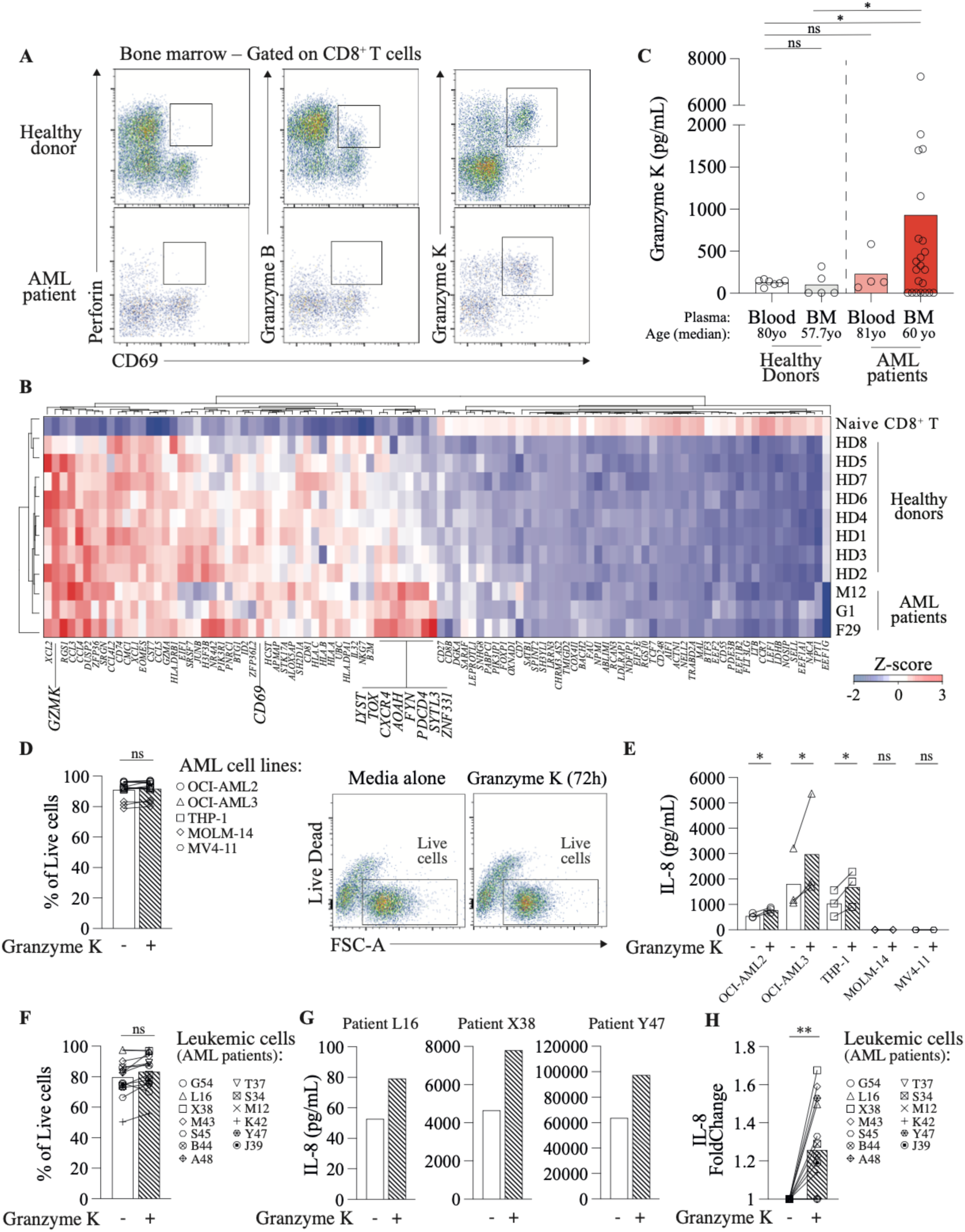
Pro-inflammatory role of CD8^+^ T cells producing Granzyme K in AML. **A.** Mass cytometry dot plots illustrating the expression of Perforin, Granzyme B, and K in relation to CD69 in healthy donor (upper panel) and AML patient (bottom panel). Data representative of one donor and one patient. **B.** Heatmap showing Z-score normalizes average expression of the top 100 differentially expressed genes identified by CITE-seq in CD69^+^ CD8^+^ T cells from the bone marrow of healthy donors (n=8) and AML patients (n=3). **C.** Quantification of Granzyme K concentrations measured by ELISA in the peripheral blood of healthy donors (n= 7) and AML patients (n= 4), and in medullary plasma of healthy donors (n= 5) and AML patients (n= 23). Statistical testing was done by unpaired t-test with Welch’s correction, two-tailed. *p ≤ 0.05, **p ≤ 0.01, ***p ≤ 0.001. **D.** Quantification of the cell viability of AML cells lines (OCI-AML2, OCI-AML3, THP-1, MOLM-14, MV4-11) after 3 days of culture, either with media alone (blank) or with Granzyme K (black stripes) (left panel) and the representative flow cytometry dot plot showing live/dead staining for AML cell line under media alone or Granzyme K (left panel). Statistical testing was done by paired t-test, two-tailed. *p ≤ 0.05, **p ≤ 0.01, ***p ≤ 0.001. **E.** IL-8 production was measured by after 3 days of culture, either with media alone (blank) or with Granzyme K (black stripes) in AML cell lines. Statistical testing was done by paired t-test, two-tailed. *p ≤ 0.05, **p ≤ 0.01, ***p ≤ 0.001. **F.** Quantification of the cell viability of patient’s leukemic cells after 3 days of culture, either with media alone (blank) or with Granzyme K (black stripes) (n=13). Statistical testing was done by Mann-Whitney test, two-tailed. *p ≤ 0.05, **p ≤ 0.01, ***p ≤ 0.001. **G.** IL-8 production by leukemic cells was measured by after 3 days of culture, either with media alone (blank) or with Granzyme K (black stripes) in AML patients. **H.** IL-8 foldchange between media and stimulation with Granzyme K was assessed on AML patients (n=13). Statistical testing was done by paired t-test, two-tailed. *p ≤ 0.05, **p ≤ 0.01, ***p ≤ 0.001.

## Discussion

Our findings reveal that CD8^+^ T cells within the AML bone marrow constitute a highly heterogeneous population, both within the tissue and across patients. Unlike in solid tumors, CD8^+^ T cells in AML bone marrow do not exhibit classical signs of exhaustion, a hallmark of chronically stimulated tumor-specific T cells. Although PD-1 expression is detected, these cells remain functional and do not co-express multiple inhibitory receptors, such as CD39 or Tim-3, suggesting that T cell exhaustion does not play a central role in AML at diagnosis^11^. Of note, similar observation has been recently reported in others hematological malignancy, such as multiple myeloma^20^. The presence of naïve, effector, and Temra CD8^+^ T cells populations lacking tissue residency markers (CD69 or CD103), coupled with their phenotypic, transcriptomic, and TCR repertoire similarities to peripheral blood CD8^+^ T cells, suggests that these cells are not intrinsic to the bone marrow microenvironment. Instead, they likely represent peripheral blood contamination during bone marrow aspiration leading to hemodilution ^22–24,27,28^. Alternatively, CD69^-^ CD8^+^ T cells may be continuously recirculating between the blood and bone marrow. However, this mechanism is primarily linked to memory T-cell maintenance and does not fully explain the presence of naïve T cells^29^. Additionally, we did not observe phenotypic or transcriptomic alterations in chemokine receptors or adhesion molecules that would suggest an active homing mechanism for these cells^30^. Finally, while tertiary lymphoid structures have been identified in solid tumors^31^, and could explain the presence of naïve cells, such structures have not been reported in AML or other hematologic malignancies. Collectively, our findings suggest that CD8^+^ T cells lacking tissue residency markers in AML bone marrow likely originate from peripheral blood contamination or recirculation rather than being resident to the bone marrow microenvironment.

In contrast, we identified distinct populations of CD8^+^ T cells expressing CD69, which are specific to the bone marrow. Within this subset, we distinguish two populations based on CD103 expression. The first, a minor fraction (<10%), comprises CD69^+^ CD103^+^ CD8^+^ T cells. These cells are not exhausted, functionally competent, and align with the definition of tissue-resident memory T cells (Trm)^19^. A similar population has been reported in the bone marrow of healthy individuals^29,32^, where they provide long-term immunity by remaining in tissues and responding rapidly to previously encountered pathogens, such as Epstein–Barr virus or influenza virus. The second, larger subset (∼90%), lacks CD103 and is characterized by an expanded TCR repertoire with distinct transcriptomic and phenotypic profiles. Despite expressing PD-1, these cells do not co-express multiple inhibitory receptors and remain functionally active. Antigenic screening revealed that they are partially specific for EBV and human cytomegalovirus epitopes. However, we cannot exclude the possibility that a subset of these cells may be tumor-specific^33^. Our findings align with previous reports demonstrating the accumulation of virus-specific CD8^+^ T cells with a unique homing phenotype in the bone marrow of healthy individuals, where they serve as a reservoir for memory T cells^30^.

Interestingly, these functional CD69^+^ CD8^+^ T cells did not exhibit the conventional cytotoxic profile characterized by Granzyme B and Perforin expression. Instead, they were marked by high levels of Granzyme K. While Granzyme K is traditionally associated with cytotoxic activity, emerging evidence suggests alternative functions, including inhibition of viral replication^34^, endothelial activation^35^, complement activation^36^, and modulation of pro-inflammatory cytokine responses^37–39^. Given their presence in the bone marrow of healthy donors, we can conclude that these cells were already present in AML patients prior to disease onset. However, their activation and secretion of Granzyme K appear to be specific to the AML context. We detected elevated concentrations of Granzyme K in the medullary plasma of AML patients, reaching levels similar to those observed in the blood plasma of sepsis patients^40^, but not in age-matched healthy donors. In the context of AML, Granzyme K did not induce leukemic cell death, excluding a pro-apoptotic role. Instead, it promoted the secretion of IL-8, a pro-inflammatory cytokine. A similar effect has been reported in endothelial cells exposed to Granzyme K^41^, suggesting that this phenomenon may also occur within the AML bone marrow microenvironment. Interestingly, we did not observe the upregulation of TNF-α^40^, IL-1β^42^, or IL-6^41^, which have been linked to Granzyme K in other pathological contexts. IL-8 plays a critical role in myeloid malignancy by promoting the migration of mesenchymal stromal cells (MSCs) into the leukemic niche^43^, enhancing AML cell expansion and chemoresistance^44^, inhibiting normal hematopoiesis^45^, and correlating with poor prognosis^46^. Moreover, multiple studies have identified a pro-inflammatory bone marrow milieu marked by elevated IL-8^47–50^ with blockade of the CXCL8–CXCR2 axis showing promise as a therapeutic approach^48^.

Due to the absence of Granzyme K in the medullary plasma of approximately 30% of AML patients, and the heterogeneous response of leukemic cells to Granzyme K among AML patients, the exact role of Granzyme K in AML and its potential specificity to a particular patient subgroup remain to explore. Moreover, we did not observe a clear association between Granzyme K expression and the mutational landscape of patients. Therefore, further analysis in larger cohort is required to determine whether these observations correlate with specific clinical features or disease subtypes. Taken together, our results highlight that rather mounting an anti-tumor response, bystander CD8^+^ T cells expressing high levels of Granzyme K in the AML bone marrow, drive the expression of pro-inflammatory IL-8 cytokines having a deleterious role in the disease progression. These findings provide a rationale for exploring therapeutic strategies aimed at inhibiting pro-inflammatory CD8^+^ T cells and targeting Granzyme K activity in association with actual therapies in cancer.

## Materials and Methods

### Study design

The objective of this study was to comprehensively characterize CD8⁺ T cells in AML and determine whether tumor-specific CD8⁺ T cells targeting leukemic cells display phenotypic features comparable to those described in solid tumors. We hypothesized that AML patients harboring such tumor-specific CD8⁺ T cells would exhibit a phenotype analogous to well-characterized CD8⁺ T cells from solid cancers. The study comprised multiple cohorts, including two reference groups based on previously published studies (*i.e.,* CD8⁺ T cells from solid tumor and non-promyelocytic AML medullary plasma)^12,50^. A discovery cohort of AML patient was analyzed using high-throughput approaches to identify phenotypic and functional signatures. Participant characteristics are detailed in Table 3 - Clinical Data. Sample sizes for each experiment are indicated in the corresponding figure legends and was based to provide sufficient power for statistical analysis. Bone marrow aspirates and peripheral blood mononuclear cells were collected from patients newly diagnosed with acute myeloid leukemia following informed written consent. The use of human AML samples was approved by the appropriate institutional review boards and ethical committees (CPP 2015-08-11DC). Medullary plasma samples from healthy donors were obtained under the “HEALTHOX” protocol (CPP Tours, AFSSAPS, ID-RCB: 2016-A00571-50; ClinicalTrials.gov identifier: NCT02789839). AML medullary plasma samples were obtained from the “MIF_AML” trial (ClinicalTrials.gov identifier: NCT03918655). Healthy bone marrow donors provided written informed consent, and sample analysis was conducted in accordance with Fred Hutchinson Cancer Center IRB approval (IRB10265) (Table 3-Clinical Data).

### Cell isolation

Bone marrow and blood samples were centrifuged at 400g for 10 minutes to separate and store plasma at-70°C. The remaining cell suspensions were layered over a Ficoll density gradient and centrifuged at 400g for 15 minutes without brake. Mononuclear cells were collected from the interface, washed in PBS, and cryopreserved in anonymized cryotubes using 90% fetal bovine serum (FBS) and 10% DMSO, and stored in liquid nitrogen.

### MHC class I tetramer assay

MHC class I tetramers with UV-cleavable peptides (allele: HLA-A*01:01, A*02:01, A*03:01, A*11:01, A*24:02, B*07:02) and streptavidin were produced in-house following published protocols^51–55^. Multiplex MHC class I assays were performed using a three-metal coding scheme (r) with 10 different metal-labeled streptavidins (n), enabling up to 120 unique tetramer combinations: 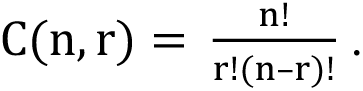 Each tetramer code was assigned a specific peptide. HLA monomers (100 μg/mL) were mixed with peptides (1 mM) in a 96-well plate, UV-exchanged at 365 nm for 10 minutes, and incubated overnight at 4°C. Tetramerization was achieved by sequential addition of streptavidin-metal conjugates according to the coding scheme, followed by incubation with free biotin (10 μM, 10 minutes). Tetramers were pooled and concentrated using a 50-kDa Amicon filter (Millipore).

### Mass cytometry sample processing

Antibodies were conjugated to metal isotopes in-house according to the manufacturer’s protocol (Standard BioTools). Cryopreserved samples were thawed and washed in staining buffer (PBS + 0.5% BSA + 0.02% sodium azide + 15 µg/mL DNase). Cells were stained with a surface antibody cocktail for 15 minutes at room temperature (RT), followed by viability staining with 5 µM cisplatin for 5 minutes, as previously described^12^. For intracellular staining, cells were permeabilized with Cyto-Fast Fix/Perm Solution (BioLegend) for 30 minutes, washed, and stained in permeabilization buffer for an additional 30 minutes. After final washes, cells were fixed in 2% paraformaldehyde (PFA) overnight. Prior to CyTOF acquisition, DNA intercalation was performed using Cell-ID Intercalator-Iridium (Standard BioTools) for 10 minutes^56^. For cytokine detection, samples were pre-stained with anti-CD69, then stimulated for 4 hours at 37°C with PMA (50 ng/mL), ionomycin (1 µg/mL), and brefeldin A (1 µg/mL) in RPMI + 10% FBS. Cells were then stained for surface and intracellular markers. For tetramer staining, cells were incubated with tetramer cocktails for 1 hour at RT, followed by surface staining^57^ (see Table 1 for antibody clones and metal conjugates).

### Mass cytometry data analysis

CyTOF data acquisition was performed as previously described^12^. CyTOF files (.fcs) were first analyzed using FlowJo to remove debris, dead cells, and contaminating cells through manual gating. CD8⁺ T cells were identified and exported into a new.fcs file for downstream analysis (Figure 1A). Subsequent analysis was conducted in R (v4.3.1). Using the flowCore package (v2.12.2), zero values were replaced with a uniform distribution between 0 and-1 to improve data visualization when using dot plot representation. Data were then transformed using the logicleTransform function (parameters: w = 0.25, t = 16,409, m = 4.5, a = 0) to approximate FlowJo scaling. Different CyTOF panels were used across patient samples, resulting in variable markers coverage (*i.e.,* n=14 to 28 patients) (Table 1 - CyTOF Antibodies Panels). To mitigate batch effects, UMAP analysis was performed on selected markers (Table 1 - CyTOF Antibodies Panels) using the umap package (v0.2.10) on transformed data, with each patient analyzed independently or grouped when staining and acquisition were performed simultaneously. Heatmaps were generated using the ComplexHeatmap package (v2.16.0), displaying median marker intensities on a color scale: black–blue for low, green–yellow for intermediate, and orange–red for high expression levels. Marker frequencies were determined by manual gating using FlowJo.

### CITE-seq sample processing

Thawed cryopreserved cells were washed in staining buffer (PBS + 0.5% BSA + 0.02% sodium azide + 15 µg/mL DNase) to remove DMSO, then stained with fluorochrome-conjugated antibodies (Table 1 - Flow Cytometry Antibodies Panel). After washing, cells were incubated with TotalSeq-C antibodies according to the manufacturer’s instructions (Table 1 - CITE-seq Antibodies Panel). Following additional washes and filtration through a 40 µm strainer, 7-AAD was added as a viability marker.

Up to 10,000 viable, flow-sorted CD8⁺ T cells were encapsulated using the 10x Chromium Controller (Supplementary Figure 2B). For each sample, libraries for gene expression, TCR sequences, and protein expression were prepared using the 5′ Chromium Next GEM Single Cell v2 Kit (10x Genomics) and sequenced on an Illumina NextSeq 500, following the manufacturer’s protocol.

### CITE-seq data analysis

FASTQ paired-end reads were processed using the Cellranger count command from 10x Genomics’ Cell Ranger software suite (v7.0.1, 10x Genomics) (Table 4 – QC sequencing) and aligned against the human reference GRCh38-1.2.0. Downstream analysis was performed in R (v4.3.1), where each patient was analyzed independently. Only single cells with sequencing information available for all three libraries (*i.e*., gene expression, TCR sequences, and protein expression) were retained for further analysis. For protein dataset visualization, integer values (n) were replaced with a uniform distribution between n and n – 1 to improve data visualization. The data were then converted to.fcs files using the flowcore package (v2.10.0). Using FlowJo software, cells positive for isotype controls from the TotalSeq-C antibody panel were excluded, as well as contaminating non-CD8^+^ T cells (CD4^+^). UMAP analysis was performed on log-transformed data using the umap package (v0.2.10) (Table 1 – CITE-seq Antibodies Panel), enabling the identification of *blood-like* cell clusters and/or CD69⁺ CD103⁺^/^⁻ CD8⁺ T cell subsets (Supplementary Figures 2B and 3). TCR repertoire analysis was conducted on single cells with sequencing information for all three libraries. Alluvial plots illustrating TCR repertoire composition were generated using the ggalluvial package (v0.12.15). Expanded TCR clonotypes were defined as groups of ≥5 cells sharing identical V(D)J segments and CDR3 sequences for paired TCR alpha and beta chains.

The frequency of expanded TCR clonotypes within *blood-like* cell clusters and CD69⁺ CD103⁺^/^⁻ CD8⁺ T cell subsets (Supplementary Figures 2B and 3) was determined using protein expression data from the same patients. Gene expression analysis of each identified expanded clonotype was performed on log-transformed data (log + 1). Differentially expressed genes (DEGs) were identified based on the highest standard deviation genes expression across expanded TCR clonotypes within each patient. Genes belonging to ribosomal protein families (*e.g.,* RPS and RPL) were excluded from the analysis and Z-scores were calculated and visualized as heatmaps using the ComplexHeatmap package (v2.16.0), where blue indicates low expression and red indicates high expression.

### Granzyme K stimulation and detection

AML cell lines (culture media: RPMI supplemented with 10% FBS and 1% L-Glutamine–Penicillin–Streptomycin) and primary leukemic cells from AML patients (culture media IMDM supplemented with 15% BIT 1% L-Glutamine– Penicillin–Streptomycin) were flow-sorted and cultured in either control medium or with recombinant Granzyme K (150 nM; Abcam, #ab157277) for 72 hours at a density of 2 × 10⁶ cells/mL. After 72 hours, culture supernatants were collected, and cytokine levels (IL-6, IL-8, IL-1β, TNFα, and S100A8/A9) were measured using ELISA kits according to the manufacturers’ protocols. Granzyme K concentrations in patient plasma samples were measured using an ELISA assay (BioTechne, #NBP3-11793).

### Statistical analysis

FACS and CyTOF data were analyzed using FlowJo software (version 10.4). Statistical analyses were performed using GraphPad Prism (version 10, GraphPad Software). Normality of the data was assessed using the Shapiro–Wilk test. Depending on the distribution, comparisons between groups were performed using either paired and unpaired Student’s t test or Mann–Whitney U tests. When unpaired two-tailed Student’s *t*-tests were used, equality of variances was assessed by F-test, and Welch’s correction was applied when variances were unequal.

## Supporting information

Table 1- Antibodies list

Table 2-Peptide list

Table 3-Clinical data

Table 4-QC CITEseq

## Supplementary Materials

Fig S1 to S9 for multiple supplementary figures

Tables 1 to 4 for multiple supplementary tables

## Data availability

Solid Cancer mass-cytometry data are available on flow-repository website: http://flowrepository.org/id/FR-FCM-ZYWM.

AML mass-cytometry data are available on https://doi.org/10.5281/zenodo.16793897 AML CITE-seq data are accessible through GEO Series accession number GSE305118.

## Authorship Contributions

Conceptualization, L.A., Y.S.; Methodology, L.A. N.D., A.A., K.M., Z.F.D., A.B., Y.S.; Data curation, L.A., N.D., I.S., R.V., C.F., K.M., M.V., J.D., O.K., F.D., O.H., R.B., D.B., M.F., N.C., Y.S.; Formal analysis, L.A., Y.S., C.G.S.; Writing – Original Draft, L.A., Y.S.; Writing – Review & Editing, L.A., N.D., C.C., P.R., D.S-B., E.W.N., E.S., F.D., O.H., E.T., Resources, I.S., O.K., F.D., O.H., R.B., D.B., M.F., N.C.; Funding, E.T., Y.S., M.F.; All authors provided critical feedback on the manuscript.

## Supplementary Figures

**Supplementary Figure 1.**
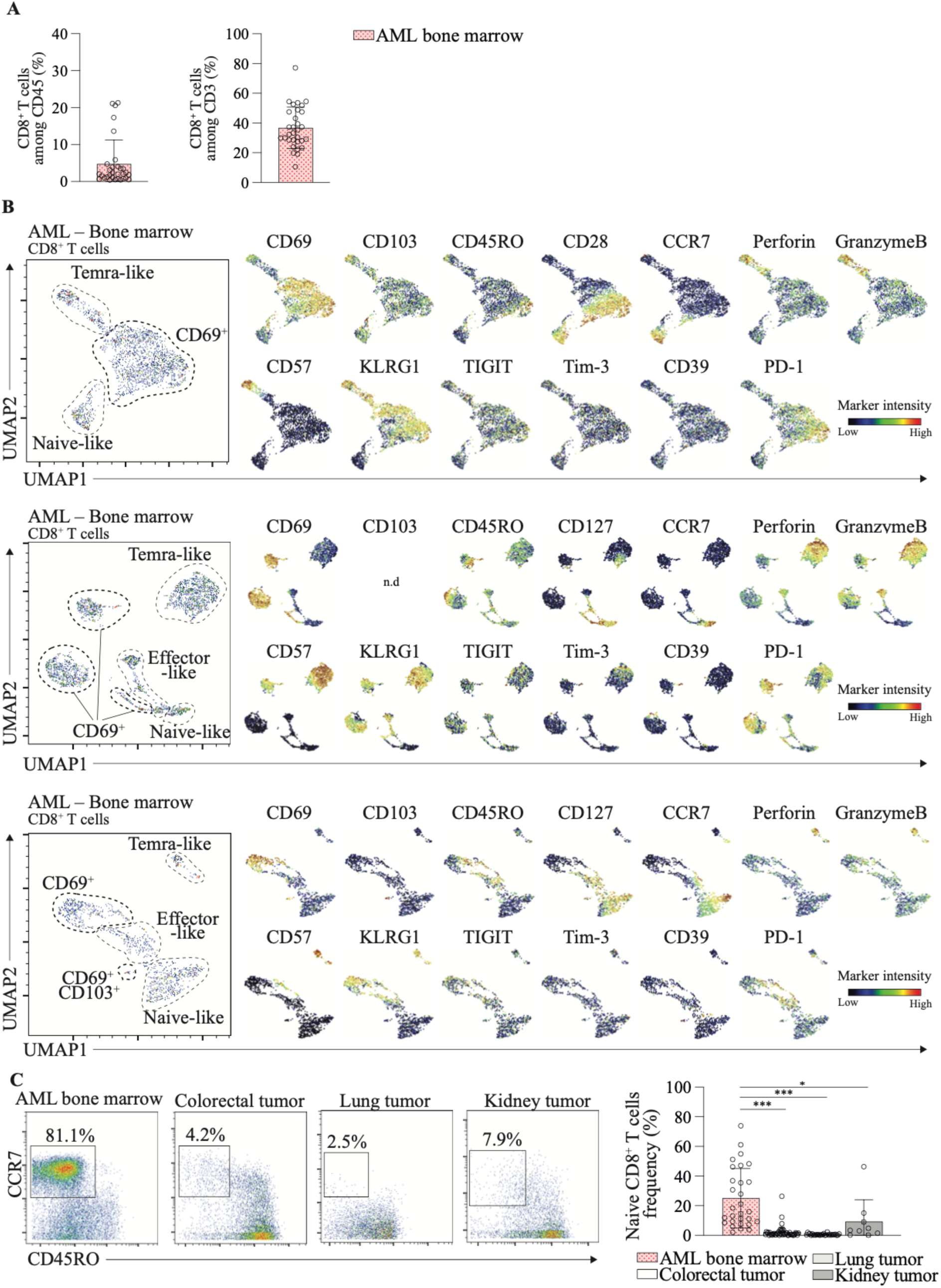
**A.** Histogram showing the proportion of CD8^+^ T cells among total CD45^+^ cells (left) and among CD3^+^ T cells (right) in bone marrow aspirates from AML patients (n=31). Data are presented as mean ± SD. **B.** UMAP plot of CD8⁺ T cells from bone marrow aspirates of an AML patient analyzed by mass cytometry (left panel). Normalized marker expression intensities were overlaid on the UMAP plot, enabling the identification of naïve-like, effector-like, Temra-like, CD69⁺, and CD69⁺ CD103⁺ CD8⁺ T cell subsets (right panel). Representative data from three patients. Mass cytometry analysis. **C.** Mass cytometry dot plot showing CD45RO and CCR7 expression on CD8⁺ T cells (left panel). Frequency of naïve CD8⁺ T cells (CCR7⁺ CD45RO⁻) in AML bone marrow (n = 29), colorectal tumors (n = 47), lung tumors (n = 29), and kidney tumors (n = 9) (right panel). Data are from biologically independent individuals. Mean ± SD; unpaired two-tailed t-test. *p ≤ 0.05, **p ≤ 0.01, ***p ≤ 0.001.

**Supplementary Figure 2.**
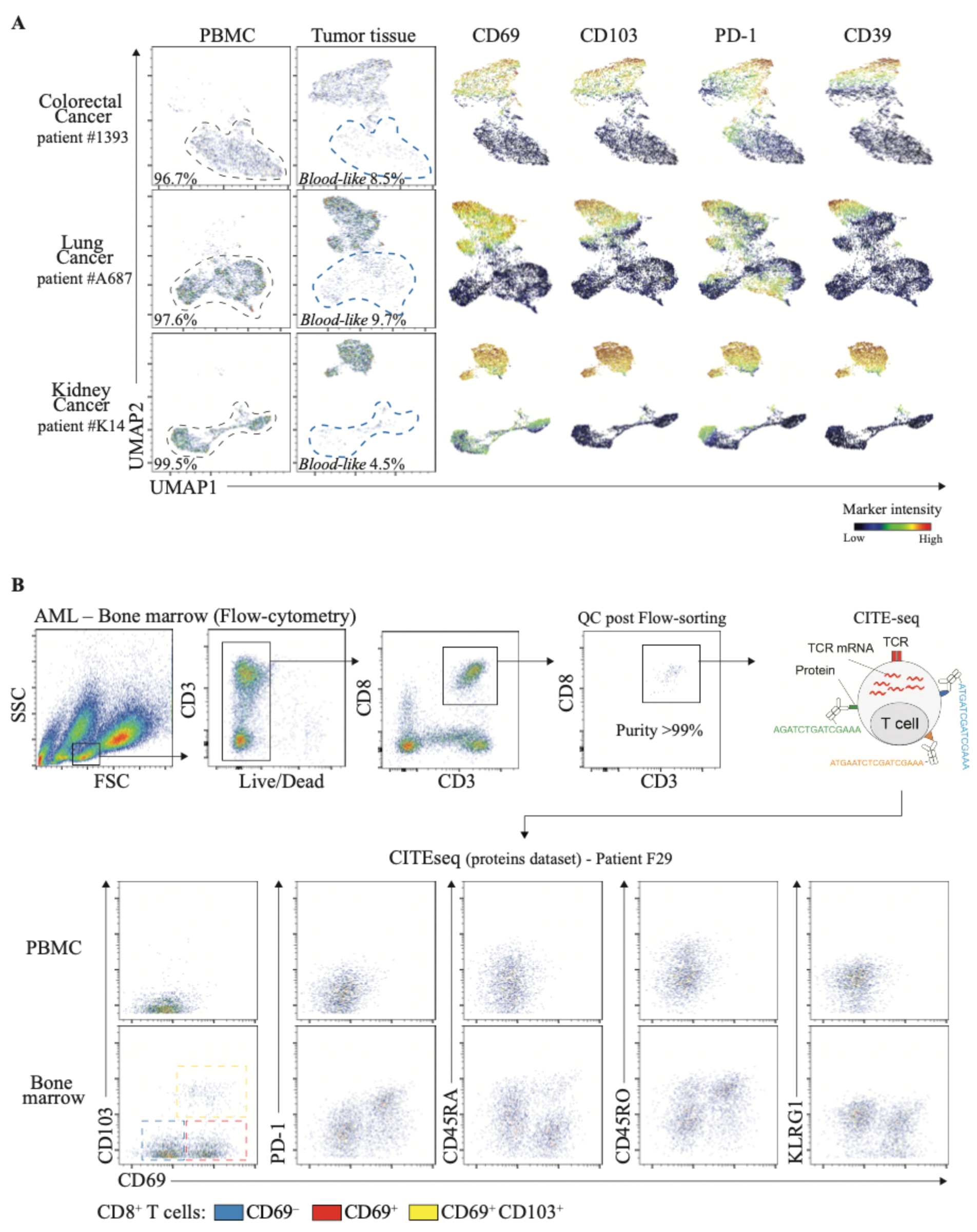
**A.** UMAP plot of CD8⁺ T cells from PBMC and tumor tissues (colorectal, lung, and kidney cancers), analyzed by mass cytometry (left panel). Normalized expression intensities of CD69, CD103, PD-1, and CD39 were overlaid on the UMAP plot (combined PBMC and tumor tissues, right panel). Representative data from three patients (see Figure 2C). **B.** Schematic representation of the CITE-seq (Cellular Indexing of Transcriptomes and Epitopes by Sequencing) experimental design for the analysis of CD8⁺ T cells. Representative data from one patient.

**Supplementary Figure 3.**
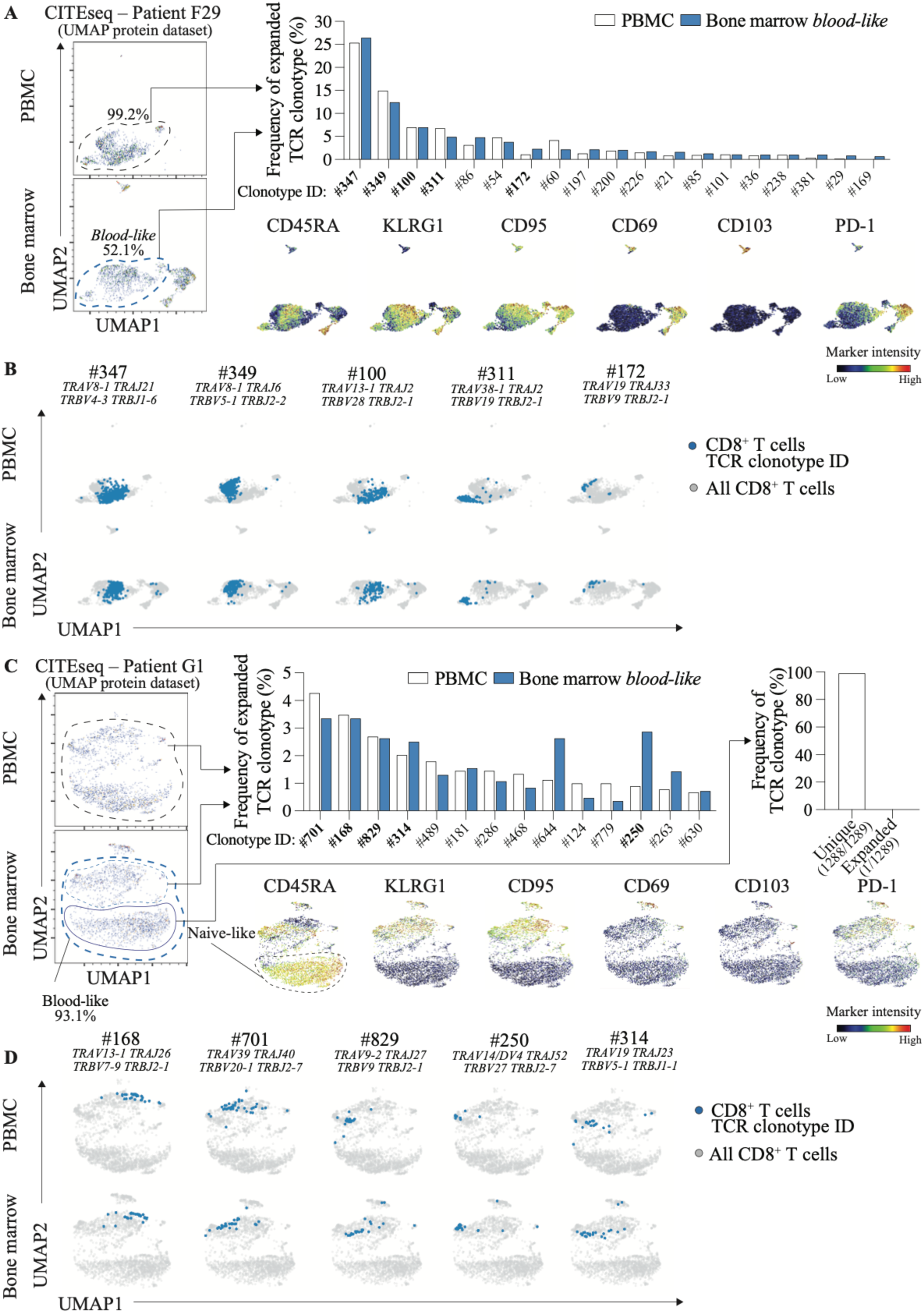
**A.** UMAP plot of CD8⁺ T cells from PBMC and bone marrow of AML patient F29, analyzed by CITE-seq. UMAP was generated using surface protein expression data (left panel). Normalized expression intensities of CD45RA, KLRG1, CD95, CD69, CD103, and PD-1 were overlaid on the UMAP (combined PBMC and bone marrow, bottom panels). The frequency of expanded TCR clonotypes among CD8⁺ T cells from PBMC and bone marrow-derived *blood-like* cells is shown (right panel). **B.** UMAP plot of CD8⁺ T cells from PBMC and bone marrow of AML patient F29, analyzed by CITE-seq. Overrepresented expanded TCR clonotypes (defined as ≥5 cells sharing identical paired V(D)J sequences) are mapped and overlaid onto the UMAP. **C.** UMAP plot of CD8⁺ T cells from PBMC and bone marrow of AML patient G1, analyzed by CITE-seq. UMAP was generated using surface protein expression data (left panel). Normalized expression intensities of CD45RA, KLRG1, CD95, CD69, CD103, and PD-1 were overlaid on the UMAP (combined PBMC and bone marrow, bottom panels). The frequency of expanded TCR clonotypes among CD8⁺ T cells from PBMC and bone marrow-derived *blood-like* cells is shown (right panel). **D.** UMAP plot of CD8⁺ T cells from PBMC and bone marrow of AML patient G1, analyzed by CITE-seq. Overrepresented expanded TCR clonotypes (defined as ≥5 cells sharing identical paired V(D)J sequences) are mapped and overlaid onto the UMAP.

**Supplementary Figure 4.**
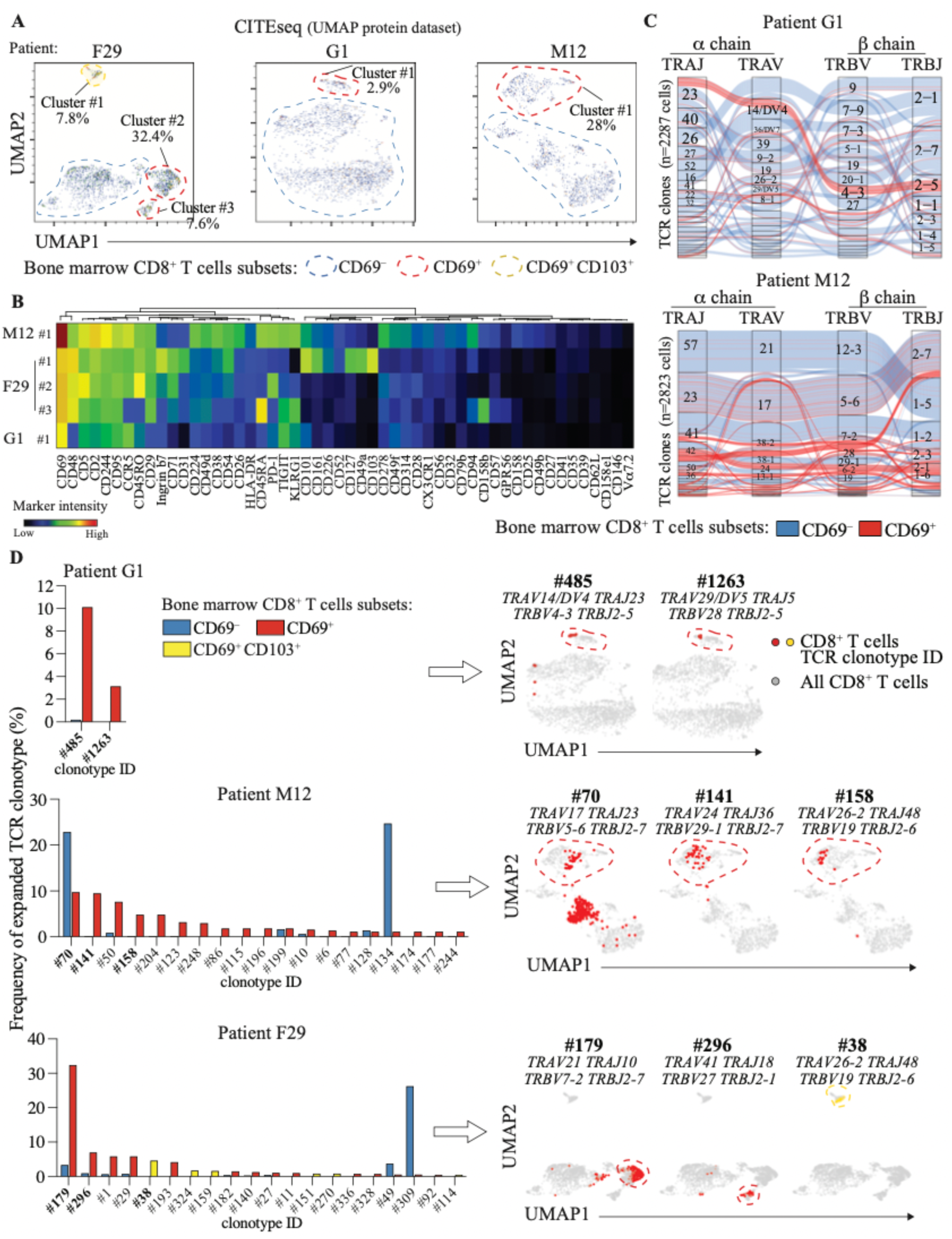
**A.** UMAP plot of CD8⁺ T cells from PBMC and bone marrow of AML patients F29, G1, and M12, analyzed by CITE-seq. The UMAP was generated using surface protein expression data (left panel). **B.** Heatmap showing median surface protein expression levels of CD8⁺ T cells from the bone marrow, comparing CD69⁺ and CD69⁺CD103⁺ clusters. Data are derived from three AML patients (G1, F29, and M12), with patient F29 exhibiting three distinct bone marrow-specific clusters. **C.** Alluvial plot illustrating the TCR clonotype composition, defined by unique paired V(D)J nucleotide sequences of the α and β chains, of CD69⁻ (blue) and CD69⁺ (red) CD8⁺ T cells from the bone marrow of AML patients G1 and M12, as measured by CITE-seq. **D.** Frequency of expanded TCR clonotypes (defined as ≥5 cells sharing identical paired V(D)J sequences) from bone marrow CD8⁺ T cells analyzed by CITE-seq (left panel). UMAP overlays display the spatial distribution of representative expanded TCR clonotypes (right panel).

**Supplementary Figure 5.**
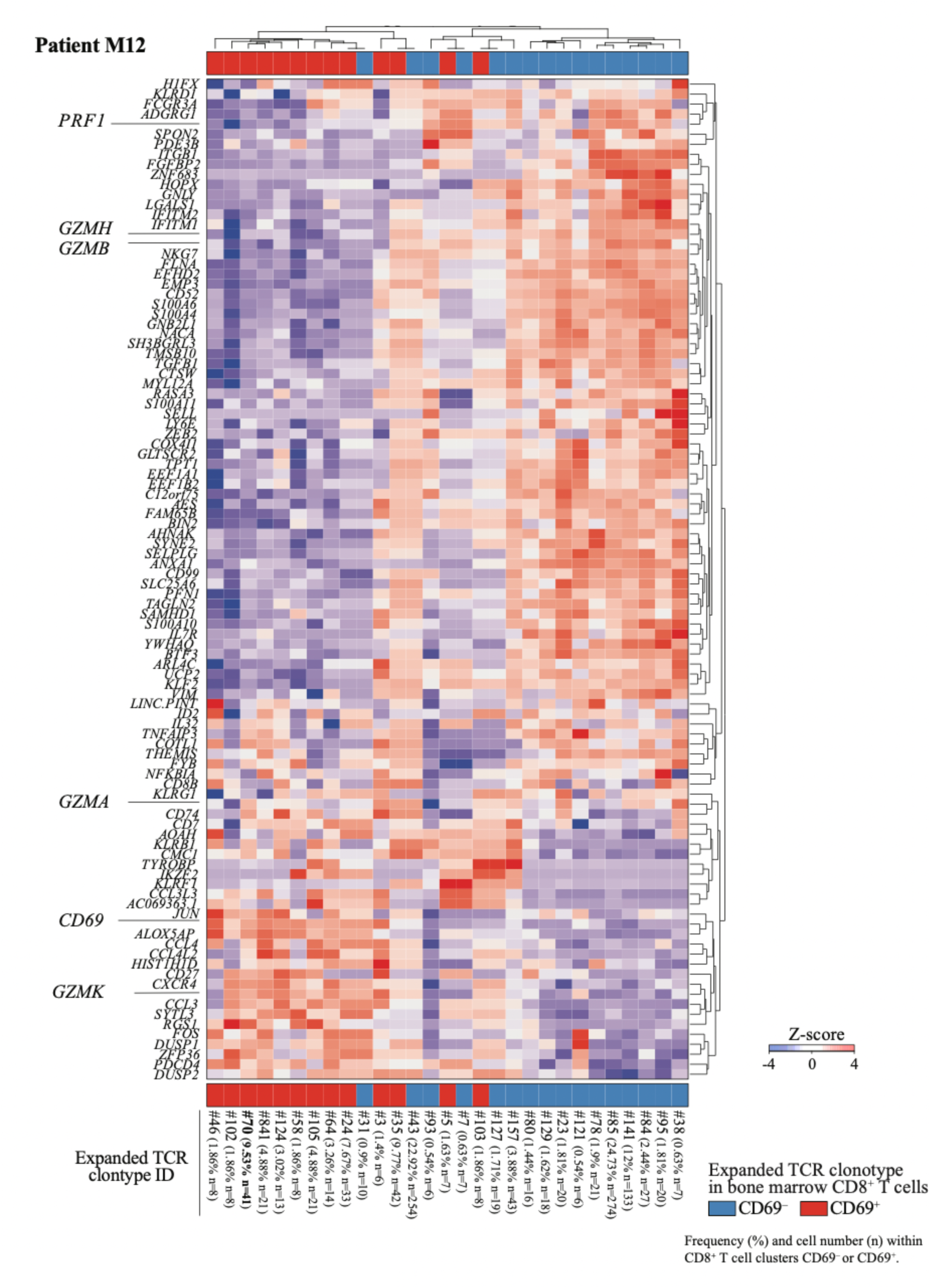
Heatmap displaying differentially expressed genes associated with expanded TCR clonotypes in CD69⁺ and CD69⁻ CD8⁺ T cells from the bone marrow of AML patient M12, as measured by CITE-seq.

**Supplementary Figure 6.**
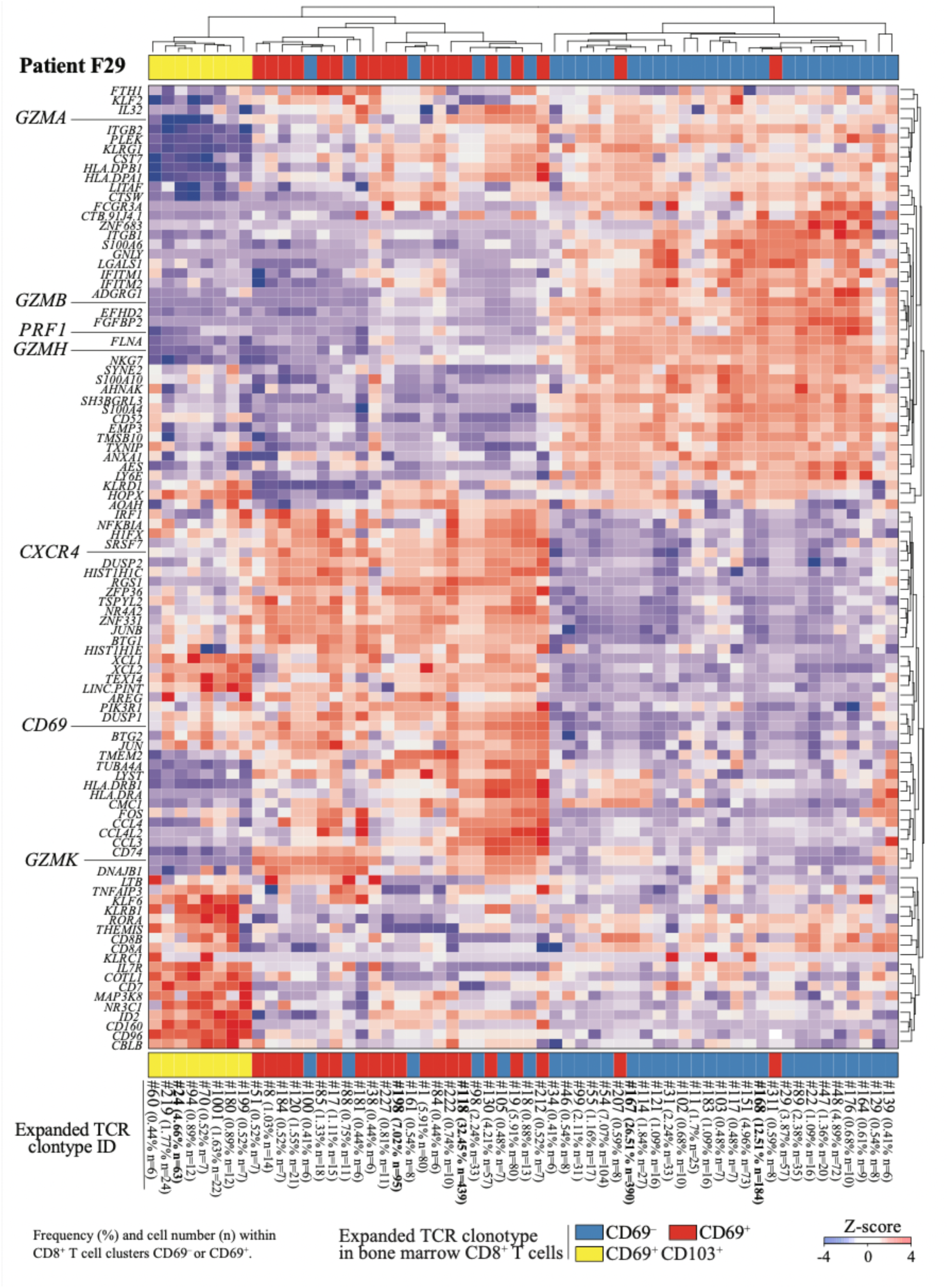
Heatmap displaying differentially expressed genes associated with expanded TCR clonotypes in CD69⁺ and CD69⁻ CD8⁺ T cells from the bone marrow of AML patient F29, as measured by CITE-seq.

**Supplementary Figure 7.**
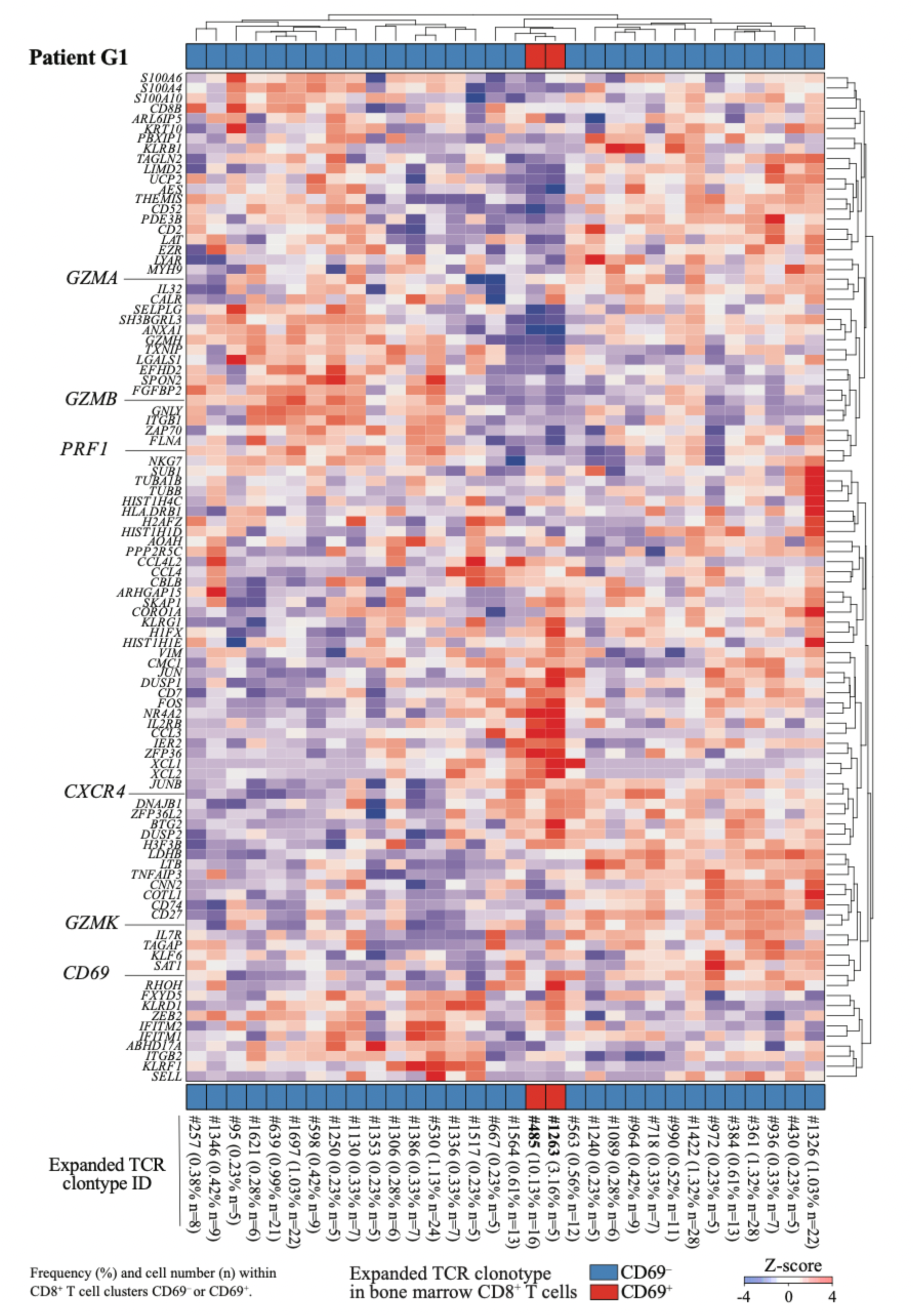
Heatmap displaying differentially expressed genes associated with expanded TCR clonotypes in CD69⁺ and CD69⁻ CD8⁺ T cells from the bone marrow of AML patient G1, as measured by CITE-seq.

**Supplementary Figure 8.**
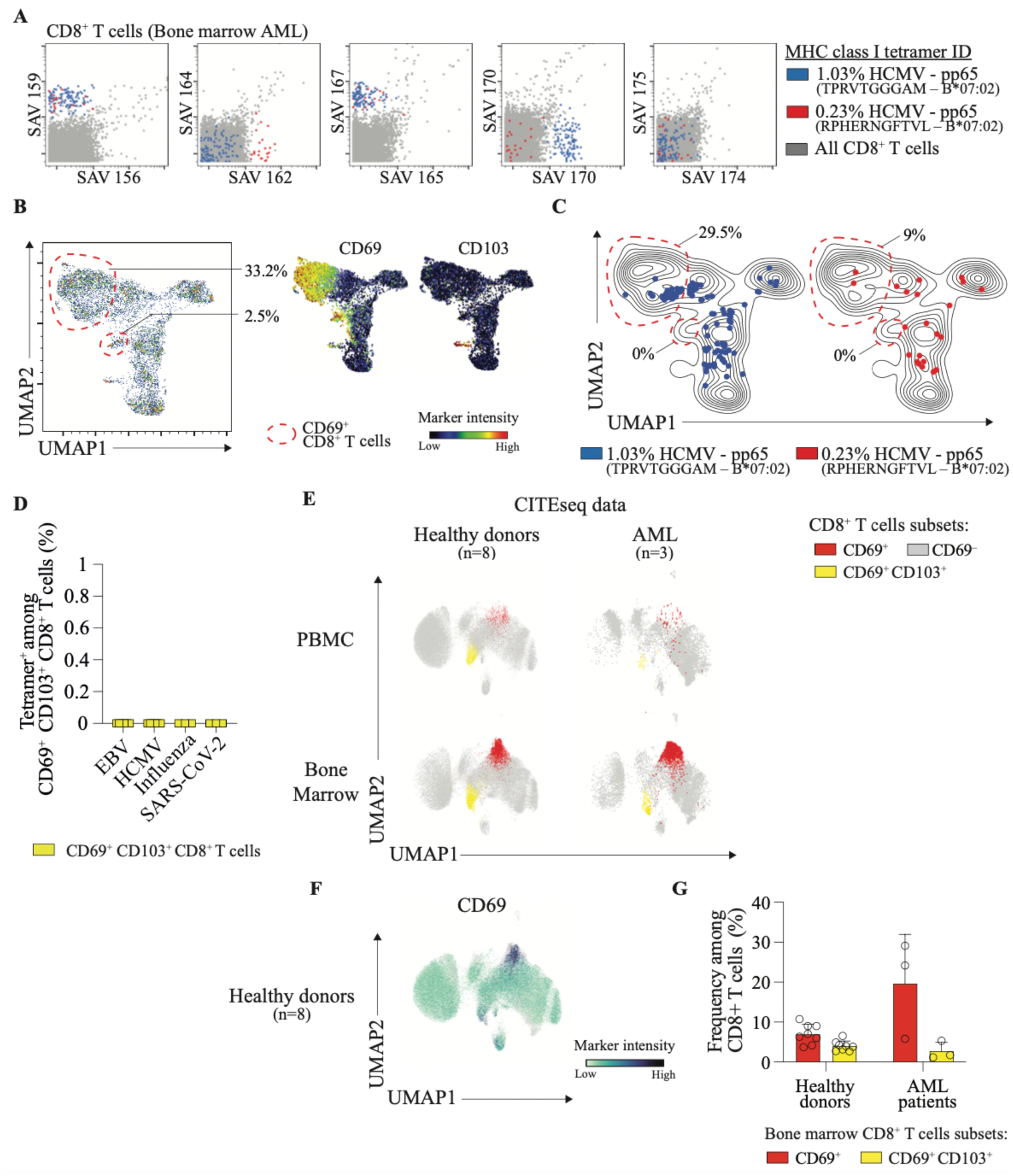
**A.** To investigate the antigen specificity of CD8⁺ in bone marrow aspirates, we performed multiplex MHC-tetramer staining as previously described (see Methods). Using a three-metal coding scheme, we simultaneously analyzed up to 120 distinct MHC tetramers targeting cancer-unrelated epitopes from EBV, HCMV, Influenza and Sars-Cov2. The complete list of peptides is provided in Table 2. Data were obtained from at least 3 independent mass cytometry experiments (see Figure 4). Representative results are shown for one patient. Each MHC tetramer-positive cell population is uniquely identified by a specific combination of three differently labeled MHC-streptavidin molecules. **B.** UMAP plot of CD8⁺ T cells from AML bone marrow analyzed by mass cytometry. Normalized expression intensities of CD69, CD103 were overlaid on the UMAP plot. Representative data from one patient. **C.** UMAP overlays show the spatial distribution of HCMV-specific CD8⁺ T cell populations identified using the MHC class I tetramer approach (see panel A). **D**. Frequencies of cancer-unrelated CD8^+^ T cells identified by mass cytometry in CD69^+^ CD103^+^ CD8^+^ T cells (yellow) from bone marrow using multiplex MHC-tetramer. **E.** UMAP plot of CD8⁺ T cells from bone marrow aspirates and blood of AML patients (n=3) and healthy donors (n=8) analyzed by CITEseq displaying CD69^-^ (grey) CD69^+^ (red) CD69^+^ CD103^+^ (yellow) CD8^+^ T cells. **F.** UMAP plot of average CD69 expression in bone marrow CD8^+^ T cells from healthy donors (n=8). **G**. Quantifications of CD69^+^ (red) CD69^+^ CD103^+^ (yellow) CD8^+^ T cells in the bone marrow of healthy donors (n=8) and AML patients (n=3), analyzed by CITEseq.

**Supplementary Figure 9.**
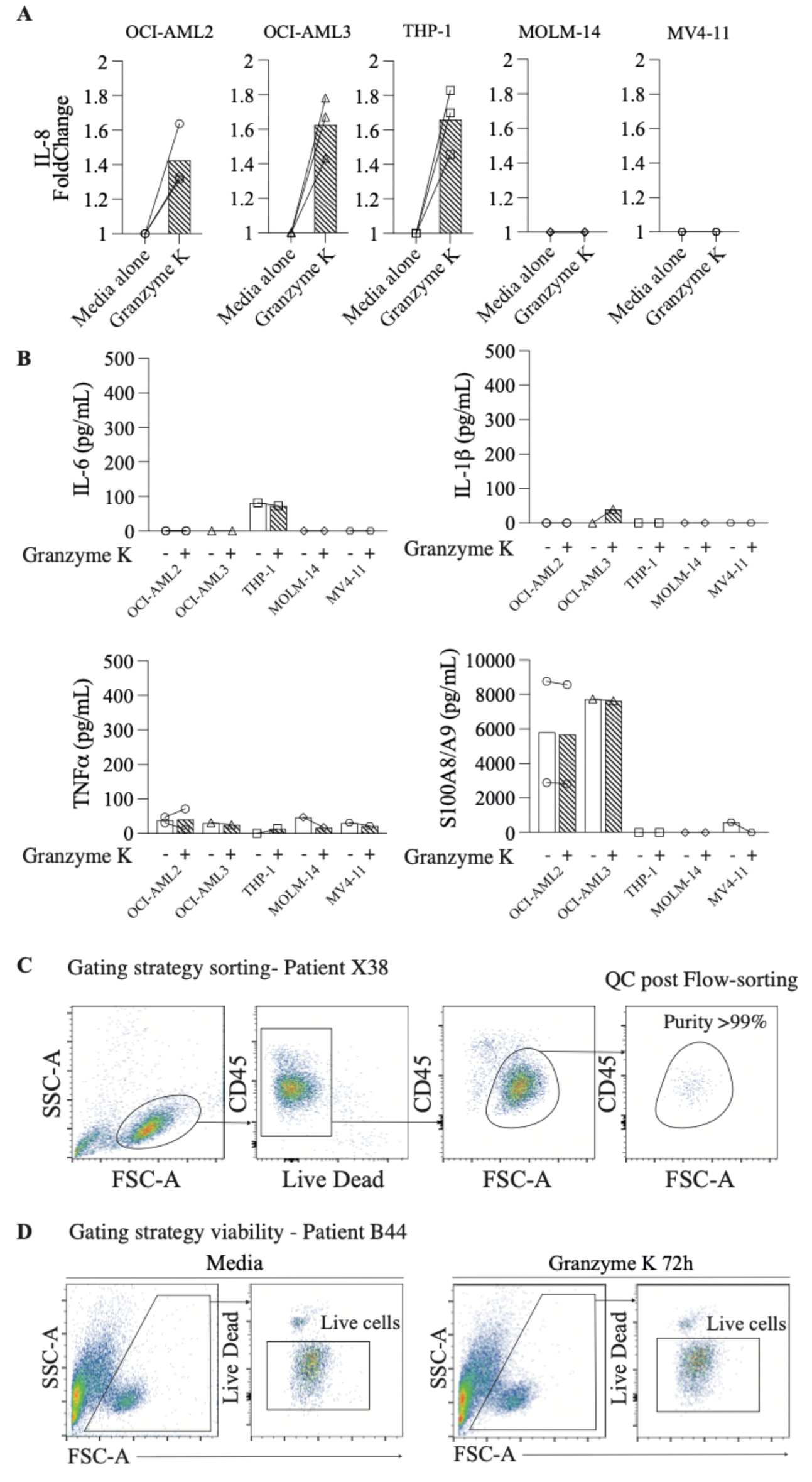
**A.** IL-8 foldchange between media and stimulation with Granzyme K was assessed on AML cell lines; OCI-AML2, OCI-AML3, THP-1, MOLM-14 and MV4-11, after 3 days of culture. **B.** IL-6, IL-1β, TNFα and S100A8/A9 production was measured by after 3 days of culture, either with media alone (blank) or with Granzyme K (black stripes) in AML cell lines. **C.** Gating strategy used to sort AML leukemic cells from blood or bone marrow from AML patients (gated on CD45^dim^ FSC-A^+^). Representative data from one patient. Flow cytometry analysis. **D.** Gating strategy used to assess the viability of live cells (gated on FSC-A^+^ Live Dead^-^). Representative data from one patient. Flow cytometry analysis.

